# Developmental noise and phenotypic plasticity are correlated in *Drosophila simulans*

**DOI:** 10.1101/2023.07.28.550919

**Authors:** Keita Saito, Masahito Tsuboi, Yuma Takahashi

## Abstract

Non-genetic variation is the phenotypic variation induced by the differential expression of a genotype in response to varying environmental cues and is broadly categorized into two types: phenotypic plasticity and developmental noise. These variation aspects have been suggested to play an important role in adaptive evolution; however, the mechanisms by which these two types of non-genetic variations influence the evolutionary process are currently poorly understood. Using a machine-learning based phenotyping tool, we independently quantified the phenotypic plasticity and developmental noise in the wing morphological traits of a fruit fly *Drosophila simulans.* Utilizing a rearing experiment, we demonstrated plastic responses in both wing size and shape as well as non-zero heritability of both phenotypic plasticity and developmental noise, which suggests that adaptive phenotypic plasticity can evolve via genetic accommodation in the wing morphology of *D. simulans*. We found a positive correlation between phenotypic plasticity and developmental noise, while the correlation between the plastic response to three kinds of environmental factors that were examined (nutrient condition, temperature, and light–dark cycle) were poor. These results suggest that phenotypic plasticity and developmental noise contribute to evolvability in a similar manner, however, the mechanisms that underlie the correspondence between these two variation types remains to be elucidated.

**Lay Summary:** Non-genetic variations consist of phenotypic plasticity and developmental noise, and these variations have been suggested to influence the direction and the rate of evolution. However, the role of phenotypic plasticity and developmental noise in evolutionary process is still poorly understood. Using a rearing experiment, we examined the heritability of plasticity and developmental noise, the correlation of the strength of plastic response to three kinds of environmental factors, and the relationship between plasticity and developmental noise in wing size and wing shape in *Drosophila simulans*. We found that the degree of phenotypic plasticity and developmental noise were heritable, and positively correlated with each other. Our results suggest that there two non-genetic variations dependently affect the direction and the rate of evolution together.

## Introduction

Genetic variation is the raw material of evolution. The pattern of genetic variation shapes the evolutionary trajectory of a relevant population and has been the subject of extensive research in evolutionary and ecological genetics (Arnold, 1992; Careau et al., 2015; Lande, 1979; Schluter, 1996). Historically, the fraction of phenotypic variation that is not attributable to genetic differences among individuals (e.g., non-genetic variation) has been considered to play only a minor role in evolution. However, recent studies have suggested that non-genetic variation has an important function in the evolutionary process (Danchin, 2013; Draghi, 2020; Ghalambor et al., 2007; Price et al., 2003). For instance, the “plasticity-first” hypothesis theorizes that non-genetic variation may take the lead in adaptive evolution (Fusco & Minelli, 2010; Pigliucci et al., 2006). Non-genetic variation is caused by two putatively independent sources of variation: phenotypic plasticity and developmental noise. The former is defined as an adaptive or maladaptive response to environmental variation (Bradshaw, 1965; Scheiner, 1993).

Phenotypic plasticity is the reaction norm to environmental changes, thus plasticity is caused by external factors. The latter is phenotypic variation caused by random fluctuations in chemical and physical signaling processes during development (Geiler-Samerotte et al., 2013; Kiskowski et al., 2019; Uller et al., 2018). However, unlike plasticity developmental noise depends solely on the local conditions of the developing phenotypes. Therefore, these two sources of non-genetic variation are fundamentally different in their causes, where one depends on external causes and the other on internal fluctuations. Currently, we have limited understanding on how these two kinds of variations influence the evolutionary process.

Phenotypic plasticity is suggested to guide adaptive evolution through genetic accommodation (i.e., plasticity-first evolution) (Badyaev, 2011; Levis et al., 2018; Levis & Pfennig, 2016; Pfennig et al., 2010; West-Eberhard, 2003). This process occurs when: (1) plasticity exists; (2) plasticity is heritable; (3) natural selection favors certain plastic responses, which guides the evolution of adaptive phenotypic plasticity; and (4) through this process, the pre-existing phenotypic plasticity is refined by natural selection into a functional phenotype (Levis et al., 2018). Thus, the ability of phenotypes to respond plastically and adaptively to environmental perturbations could influence the direction and rate of evolution. The rationale behind these ideas is that adaptive plasticity can first build the developmental mechanisms that generate the adaptive phenotypes more often than the non-adaptive ones (e.g., adaptive developmental bias), and these mechanisms could then later become genetically determined. To study the evolutionary significance of phenotypic plasticity via genetic accommodation, this developmental bias needs to be quantified.

One approach to quantify the developmental bias is to evaluate the developmental noise, which reflects the manner by which the developmental machinery translates the genetic and developmental inputs to the phenotypic expressions. Recent theoretical and empirical studies have suggested that the developmental noise is related to evolution (Furusawa et al., 2005; Kaneko & Furusawa, 2006; Rohner et al., 2022; K. Sato et al., 2003; Uller et al., 2018). Therefore, the two sources of non-genetic variation and evolutionary outcomes may be hierarchically related in that developmental noise represents the variability of the developmental processes which underlies the pattern of phenotypic plasticity, and evolution is predisposed to act on variations that are biased through these processes.

However, phenotypic plasticity and developmental noise are often conflated. For instance, although phenotypic plasticity and developmental noise are caused by biologically distinct processes (Scheiner et al., 1991; Scheiner, 1993), they are often lumped together as residual variation after the various types of genetic variances are estimated (Wilson et al., 2010). Moreover, the variance of developmental noise is typically measured as the difference in trait values between the left and right side of an individual (i.e., fluctuating asymmetry (FA)) (Gangestad & Thornhill, 1999; Rohner et al., 2022; Van Valen, 1962) and is notoriously difficult to evaluate. Not only does FA require repeated measurements of all the paired-traits (Palmer & Strobeck, 1986), but a reliable estimate of FA variance in a population often requires a substantial amount of samples (Houle, 1997). In this study, we overcome this logistical challenge by using a recently developed machine-learning based method to semi-automatically measure phenotypes from digital images (Porto & Voje, 2020).

Another approach to study the developmental bias is to evaluate how the different types of environmental cues induce plastic responses. The idea is that if different environmental stimuli generate phenotypic expressions through the same developmental machinery, the pattern of plasticity is expected to be similar. However, most studies on phenotypic plasticity have evaluated the reaction against only one environmental change at a time (De Lisle et al., 2022; Hudak & Dybdahl, 2023; Levis et al., 2018; Nyamukondiwa et al., 2011; A. Sato & Takahashi, 2022; Westneat et al., 2019). Nutrition (Burns, 1992; Shehata & Marr, 1971) and temperature (Yamahira et al., 2007) are two of the most commonly studied environmental stimuli on phenotypic plasticity, and are considered to influence the phenotypic expression through the variation in growth rate. Therefore, these two plastic responses might exhibit similar patterns. Another stimulus that has been of great interest in insects is light–dark cycle because it determines voltinism (Lindestad et al., 2019). Plastic responses to the light–dark cycle are most likely driven by altering the circadian rhythm, which may not be directly related to the growth rate and induces a different response pattern compared to those of nutrition and temperature.

Although the non-genetic variation is not heritable through the standard inheritance mechanism, the propensity of a genotype to respond to internal and external perturbations are heritable. For example, in *Polygonum cespitosum*, an Asian annual plant, the ability of a population to respond plastically to environmental conditions varies between the populations (Matesanz et al., 2012). Some theoretical and empirical studies have shown that the phenotypic plasticity ability is varied within a single population (Levis et al., 2018; Pigliucci et al., 2006). Similarly, the degree of developmental noise is known to vary within and between populations (Kiskowski et al., 2019). Genetic variation in the ability to produce phenotypic plasticity and developmental noise are potentially important contributors to the ability of a population to respond selectively (Hansen & Houle, 2008).

Here, we revisited the relationship between the developmental noise and phenotypic plasticity of the wing morphological traits of *Drosophila simulans*. We first estimated the phenotypic plasticity by measuring the plastic response of iso-female lines to multiple environmental conditions and developmental noise by FA. We then calculated the heritability of the phenotypic plasticity and developmental noise. Finally, we examined the correlation between the phenotypic plasticity developmental noise.

## Methods

### Study species, sampling, and rearing

*Drosophila simulans* is a common fruit fly species in Japan. We captured adult individuals of *D. simulans* from the campus of Chiba University, Japan (35° 62′ 79′′ N, 140° 10′ 31′′ E) in 2020 and established iso-female lines. Each iso-female line was reared with a standard medium that was made based on that described by Fitzpatrick et al. (2007) (500 mL H_2_O, 50 g sucrose, 50 g dry yeast, 6.5 g agar, 5.36 g KNaC_4_H_6·_4H_2_O, 0.5 g KH_2_PO_4_, 0.25 g NaCl, 0.25 g MgCl_2_, 0.25 g CaCl_2_, 0.35 g Fe_2_(SO_4_) _·_6.9H_2_O) in 170 mL bottles under a 12 h light–dark cycle at 25°C. Each iso-female line was repeatedly inbred over 20 generations to remove genetic variation in a line and the maternal effect.

### Strains and wing collection

Six iso-female lines were randomly chosen and used for the following experiments. The degree of phenotypic plasticity was quantified by comparing the wing morphology and wing size of individuals reared under seven combinations of three environmental factors. These combinations consisted of three nutrient conditions (high, intermediate, or low), three light–dark cycle conditions (10 h light/14 h dark, 12 h light/12 h dark, or 14 h light/10 h dark) and three temperature conditions (20°C, 23°C, or 26°C). In the present study, high, intermediate, and low nutrient media were composed of a 0%, 40%, and 80% reduction, respectively, of the active yeast and sucrose concentrations of the standard medium. For each environmental condition, one of the three environmental factors varied from the standard condition (12 h light/12 h dark, 23°C, intermediate nutrient). Since the standard condition was shared three times, the total number of environmental conditions was seven. For each iso-female line, 32 eggs were put into the vial and reared under each of the seven conditions from eggs to adults. Two days after the first adult emerged in a vial, all female adults were collected. The left and right wings of the females were separated from their bodies and placed directly onto a glass slide. A glass cover was placed over the wings and the cover glass and glass slide (i.e., dry mount) were glued to flatten the wings.

### Analysis of wing morphology and wing size

We took pictures of the wings with a CMOS camera (Leica MC190 HD, 10 million pixels) of the stereoscopic fluorescence microscope (Leica M165 FC) under constant light conditions where the wings are lit up by the torres stand under the glass slide. To evaluate the variation in the wing morphology, we measured x–y coordinates of 12 landmarks placed at the vein intersections of the wings (Fig. 1a) following Houle et al. (2017). To place landmarks to the images, we used the machine-learning program, “*ml-morph*” (Porto & Voje, 2020). First, we built a training set based on 125 wings that were manually landmarked, then used *ml-morph* to place the landmarks on 250 new images, which were trained by the first training set, and any erroneous landmarks were manually corrected. These landmarks data from 250 images were used to build the second training set as new teaching data. Next, we used *ml-morph*, trained by second training set, to place landmarks on 1800 images and manually corrected any errors. Finally, we built the third training set using the 1800 landmarks data, trained *ml-morph* with the third training set, and used this *ml-morph* to obtain the landmarks. The images that were used to build the training set included not only the images used in this study but also wing images of other *Drosophila* species. With these procedures, we improved the accuracy of the machine-learning program. Using training sets based on 1800 images, we automatically landmarked, then manually corrected any wing coordinates that were found to be incorrect by *ml-morph*. This procedure allowed us to repeatedly measure all specimens twice without additional manual effort. All right wings were horizontally flipped. In total, we landmarked the left and right side wings from 410 individuals (820 images).

**Figure 1.**
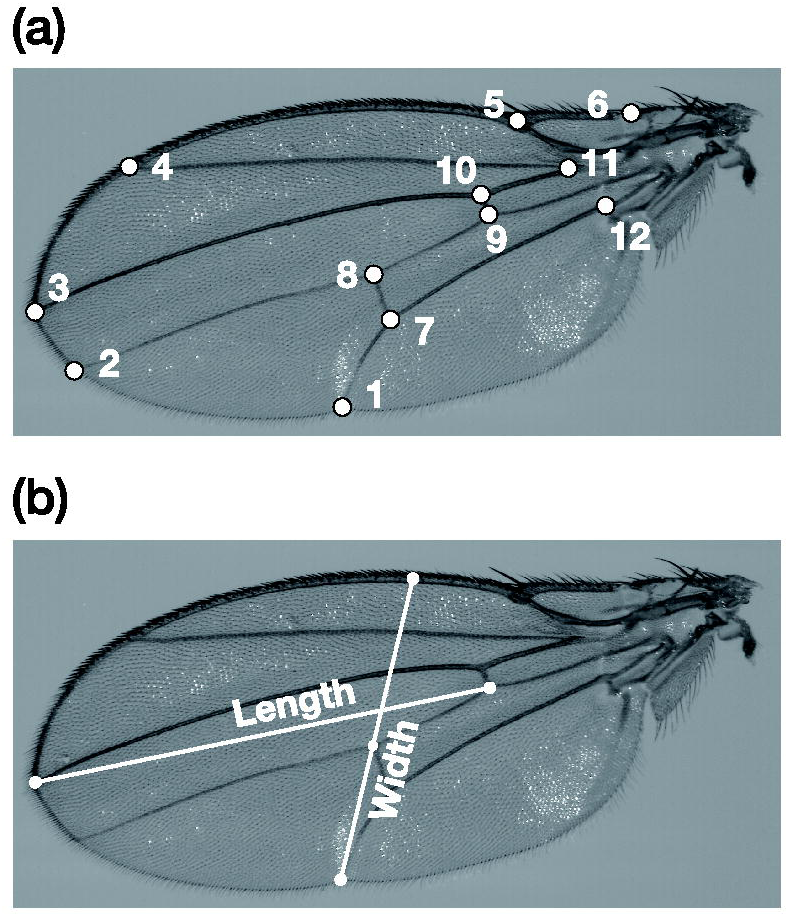
Picture of the left wing, and illustration of the wing morphology (a) and wing size (b) measurements.

All statistical analyses in this study were conducted using *R* 4.1.2. We standardized all wing coordinates using the generalized Procrustes analysis (GPA) with respect to size, rotation, and translation, which translated the original coordinates data to a common coordinate system, by using the “*geomorph*” package in *R* (Adams & Otárola-Castillo, 2013). To evaluate the variation in wing size, we also measured the wing length and width based on the method described by Lack et al. (2016) using ImageJ version 2.1.0 (https://imagej.nih.gov/ij/). For the wing length measurement, we measured a straight line drawn from the intersection of the anterior crossvein and L4 longitudinal vein, to where the L3 longitudinal vein intersects the wing margin. For the wing width, we measured a straight line from the intersection of the L5 longitudinal vein and posterior wing margin, passing through the intersection of the posterior crossvein and L4, and terminating at the anterior wing margin (Fig. 1b).

### Dimension reduction and measurement of variance

After GPA, the shape data consisted of 20 dimensions. To reduce this to a smaller number of effective morphological dimensions, we performed principal component analysis (PCA) on the averaged coordinate data, in which the original coordinates data were averaged by the iso-female lines and rearing environmental conditions. The degree of phenotypic plasticity of each iso-female line for wing morphology was defined as the standard deviation of the left wing principal component (PC) scores among the seven rearing conditions. We used the top five PCs whose contribution was greater than 5% of the total phenotypic variance (Fig. S1). The developmental noise was assessed using FA which was the difference between the right and left wing PC scores. The degree of developmental noise was then evaluated as the standard deviation of FA among individuals (Goswami et al., 2015; Klingenberg, 2019; Rohner et al., 2022). We evaluated the phenotypic plasticity and developmental noise of the wing length and wing width using the same approach.

### Correlation between plasticity under different environmental conditions

To examine the correlation among the strength of the plastic responses to each environmental factor, we used Pearson’s correlation test. For the wing morphology, we only used the left wing PC scores. PC1 to PC5 were analyzed and addressed as independent wing morphology traits. Each strength of plastic response was defined as the standard deviation of the left wing PC scores within each rearing condition. For wing size, the left wing length and wing width were used, and the strength of plastic response was evaluated in the same manner. We then applied the Pearson’s correlation test between the strength of plastic response to each environmental factor.

### Broad-sense heritability

Following a previously described method (Becker, 1964; Scheiner & Lyman, 1989), we estimated the broad-sense heritability of phenotypic plasticity as

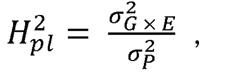

where 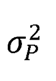 is the total phenotypic variance and 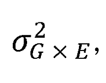 is the genotype–environment interaction. We used the first five PCs (PC1–PC5), to estimate the heritability of phenotypic plasticity in wing morphology. To estimate 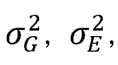, and 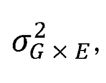, we constructed an analysis of variance (ANOVA) where the PC scores of the individual wings are the response variable, and the identity of the iso-female line, environmental condition, and their interaction term were the predictor variables. In this model, the variance components of the iso-female line, environment, and their interaction represent 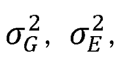, 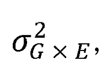, respectively. The PC scores used for the estimations were derived from the PCA performed on the standardized coordinates of the left wings. First, we calculated the heritability of each trait from its respective PC scores and then we calculated the average heritability of the five independent traits and considered them to be the heritability of the phenotype plasticity in wing morphology. Heritability of the phenotype plasticity in wing length and width were estimated in the same manner.

We defined the heritability of the developmental noise as the portion of the total FA variance that can be explained by difference among iso-female lines. The equation used is as follows:

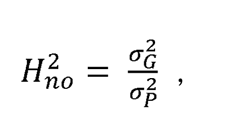

where 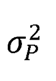 is the total FA variance among individuals and a^2^_G_ is the between-line variance of FA. To evaluate these variance components, we used ANOVA where individual FA was the response variable, and the line was the predictor variable. FA was estimated based on the difference between the right and left wing PC scores, and we used PC1–PC5 as the estimation of the heritability of plasticity. In addition, heritability of the developmental noise in wing length and width were estimated in the same manner. To estimate this heritability, we used type II ANOVA from the “*car*” package.

### Correlation between plasticity and developmental noise

To investigate the relationship between the degree of phenotypic plasticity and developmental noise, we used Pearson’s correlation test and a linear mixed-effects model from the “*lme4*” package (Bates et al., 2015). This model was fitted using the maximum likelihood with “*lmerMod*.” In the model evaluating the relationship between the wing morphological traits, we included log_10_ of plasticity as the response variable and log_10_ of developmental noise as the predictor variable and trait identity (PC1–PC5) as the random effect. The observations were weighted with the eigenvalue of each PC. In the model evaluating the relationship in wing size, we constructed the same model with trait identity (wing length or wing width) as the random effect.

## Results

### Variation in the strength of plastic response

For the wing morphology, we identified the top five PC axes whose contribution was greater than 5% (PC1: 36.3%, PC2: 26.5%, PC3: 10.6%, PC4: 7.7%, PC5: 6.3%, Fig. S1). We found a significant effect of the rearing condition on the wing morphology in PC1 but not in PC2 (Fig. 2a; PC1: *F*_6,30_ = 4.215, *P* < 0.01; PC2: *F*_6,30_ = 1.779, *P* = 0.137). The degree of plastic responses in the wing morphology along the PC2 axis were mainly dependent on the differences between lines (PC1: *F*_5,30_ = 2.316, *P* = 0.068; PC2: *F*_5,30_ = 4.853, *P* < 0.01). In contrast, both the wing length and wing width were critically dependent on the environmental conditions to which the different iso-female lines were exposed (Fig. 2b; wing length: *F*_6,921_ = 130.2, *P* < 0.001; wing width: *F*_6,921_ = 63.6, *P* < 0.001).

**Figure 2.**
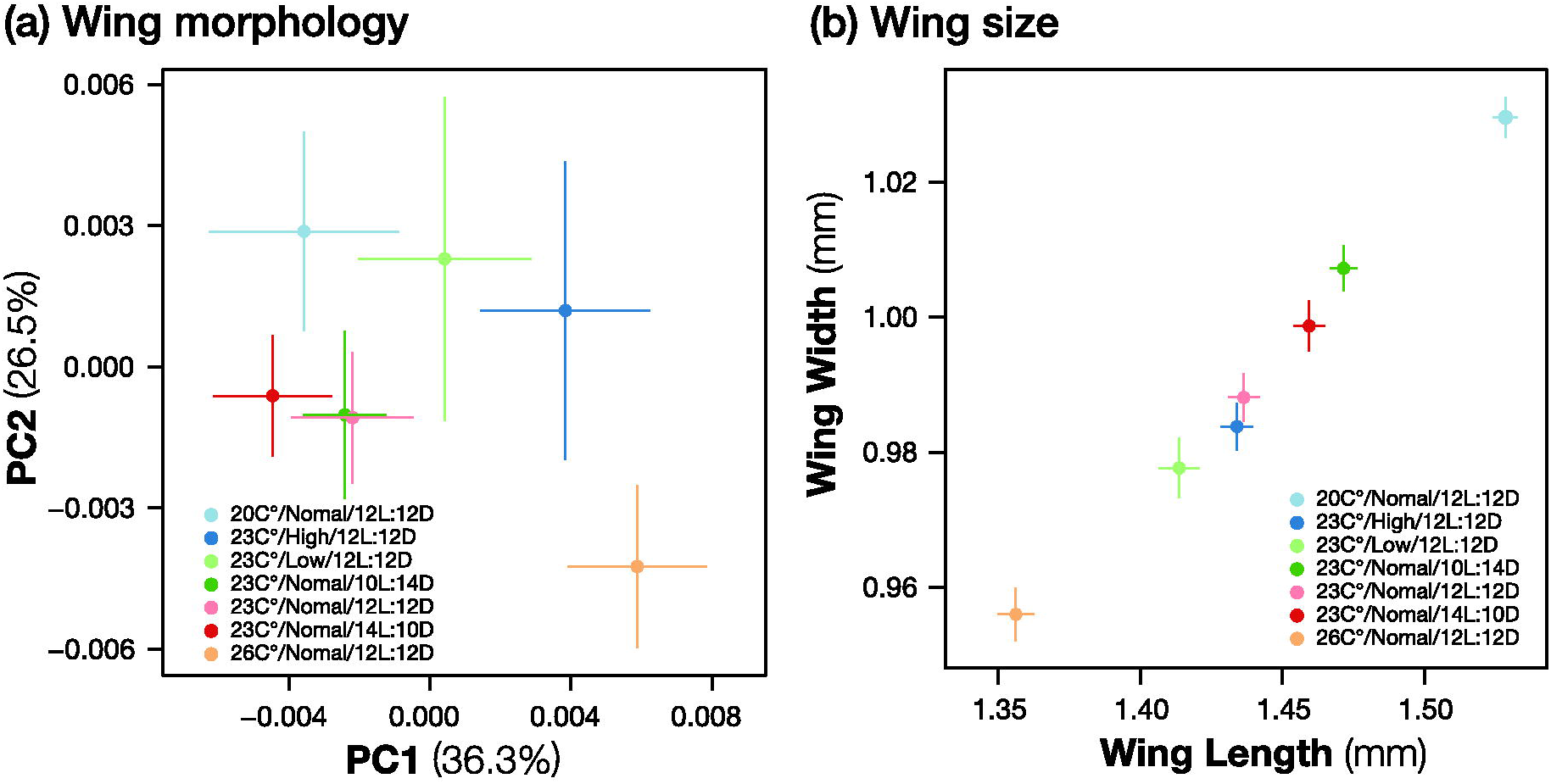
The variation in the wing morphology (a) and wing size (b) of *Drosophila simulans* among the different rearing environmental conditions. The color of points and error bars represent the environmental conditions to which the individuals were exposed. Points are the mean and error bars are the standard error of the mean.

The strength of plastic responses among the environmental factors were not always correlated (Fig. 3). Although the strength of plastic response to the nutrition and light–dark cycle conditions had significant positive correlation (*r* = 0.47, *P* < 0.01), the strength of plastic response between the temperature and nutrition conditions (*r* = 0.28, *P* = 0.14) and between the light–dark cycle and temperature conditions were not correlated (*r* = 0.04, *P* = 0.83). However, it is noteworthy that the lack of statistical significance in these analyses may partly be attributed to the small sample size and generally small effect sizes of the plasticity in wing morphological traits. A general lack of correlation between the plastic responses under different environmental conditions is further supported by our results for wing length and wing width, where no significant correlations between the strength of plastic response to each environmental factor were found (wing length: between temperature and light–dark cycle, *r* = 0.31, *P* = 0.55; between temperature and nutrition, *r* = 0.22, *P* = 0.67; between light–dark cycle and nutrition, *r* = 0.30, *p* = 0.56, wing width: between temperature and light–dark cycle, *r* = 0.10, *P* = 0.86; between temperature and nutrition, *r* = 0.57, *P* = 0.24; between light–dark cycle and nutrition, *r* = −0.25, *P* = 0.63).

**Figure 3.**
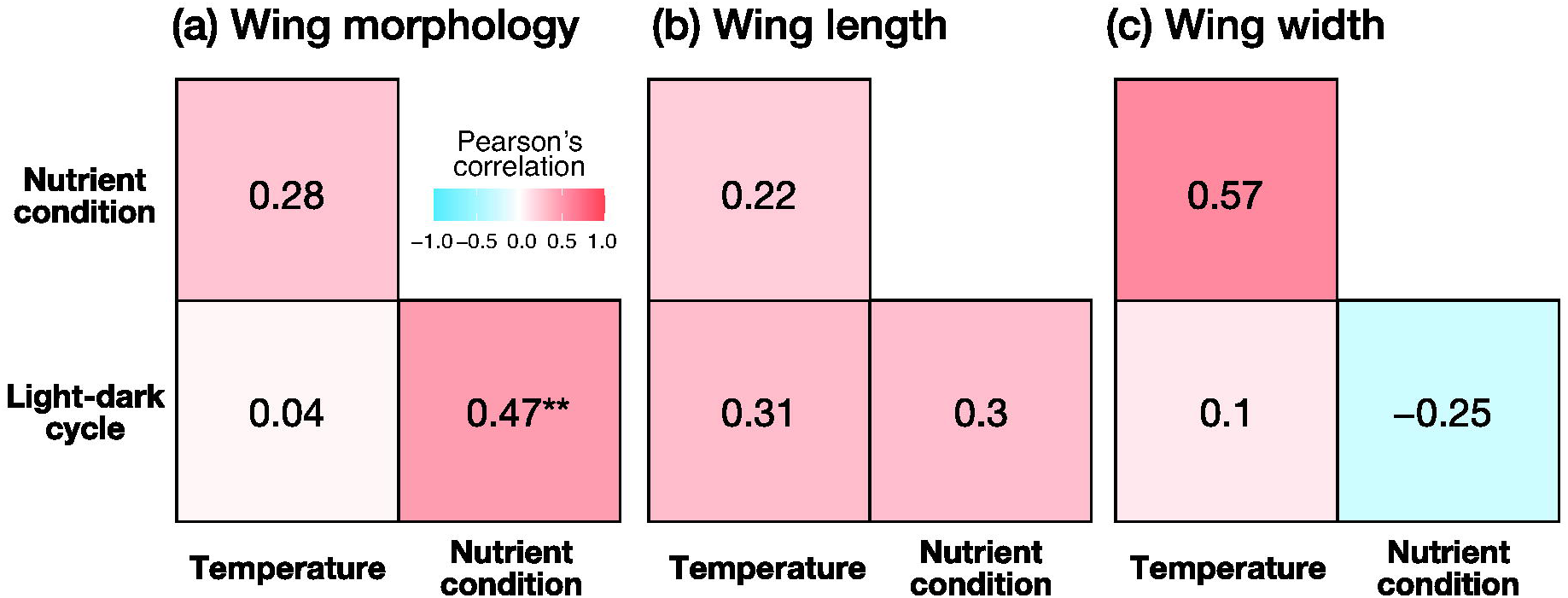
The correlation matrix heatmaps for the wing morphology (a), wing length (b), and wing width (c), respectively. The value in the center of each box represents Pearson’s correlation score. A blue box indicates negative correlation, a red box indicates positive correlation, and a white indicates no correlation.

### Broad-sense heritability of phenotypic plasticity and developmental noise

The heritability of phenotypic plasticity and developmental noise in wing morphology was 9.4% (PC1: 0.067; PC2: 0.175; PC3: 0.097; PC4: 0.091; and PC5: 0.044; mean ± standard error (*SE):* 0.094 ± 0.022) and 2.4% (PC1: 0.018; PC2: 0.044; PC3: 0.015; PC4: 0.029; and PC5: 0.011; mean ± *SE:* 0.024 ± 0.007), respectively (Fig. 4a). In addition, the heritability of the phenotypic plasticity and developmental noise in wing size was 1.4% (length: 0.023; width: 0.004; mean ± *SE:* 0.013 ± 0.010) and 0.3% (length: 0.005; width: 0.001; mean ± *SE:* 0.003 ± 0.002), respectively (Fig. 4b).

**Figure 4.**
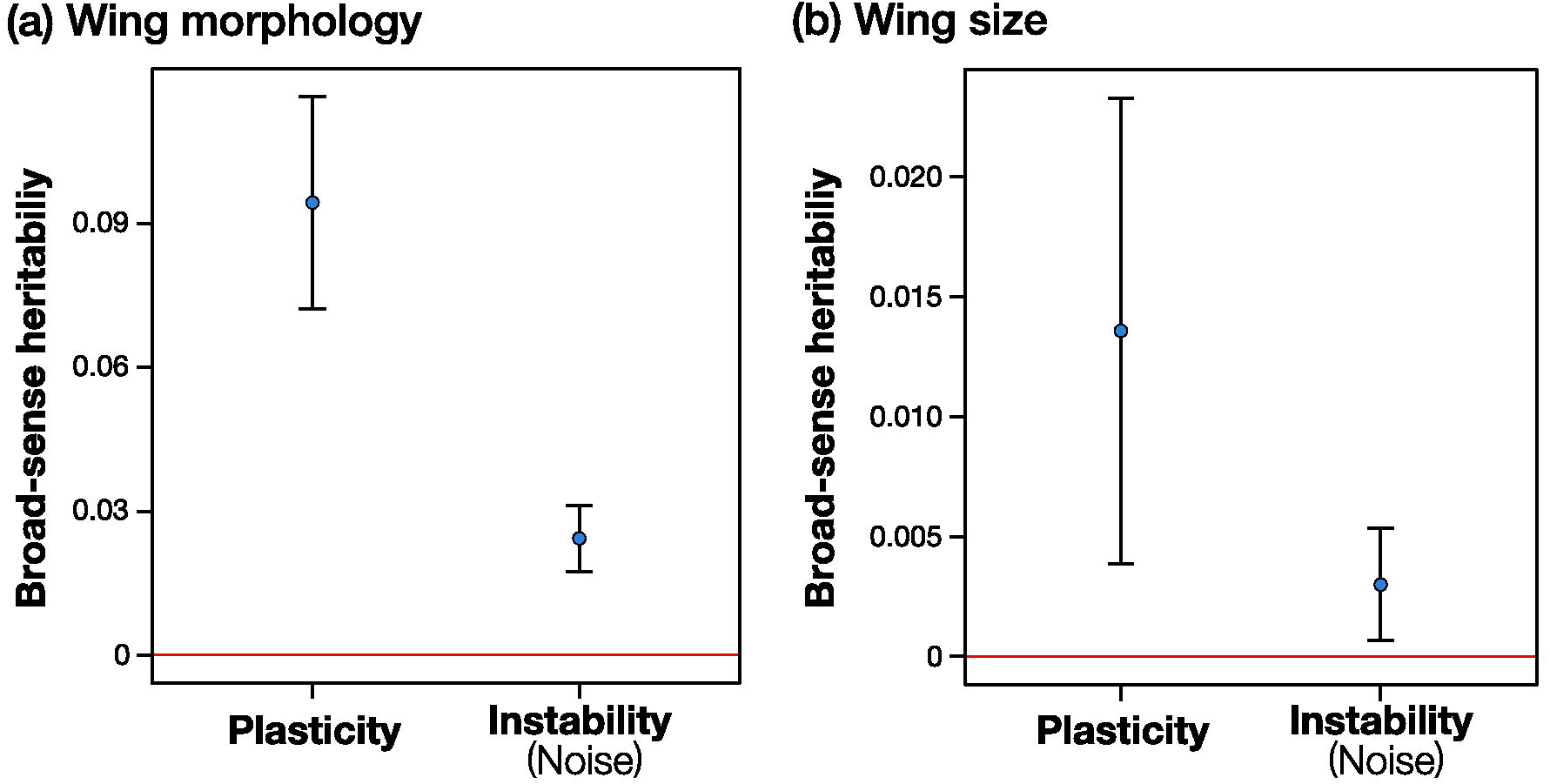
The broad-sense heritability of the phenotypic plasticity and developmental noise in the wing morphology (a) and wing size (b). Points are the mean and error bars are the standard error of the mean.

### Relationship between phenotypic plasticity and developmental noise

There was a significantly positive relationship between phenotypic plasticity and developmental noise in the wing morphology (Fig. 5a; *P* < 0.001, conditional R^2^ = 0.877). In addition, there was a positive relationship between the non-genetic variations in wing size (Fig. 5b; *P* < 0.01, conditional R^2^ = 0.773). These two types of non-genetic variations were correlated in the wing morphological traits of *D. simulans*.

**Figure 5.**
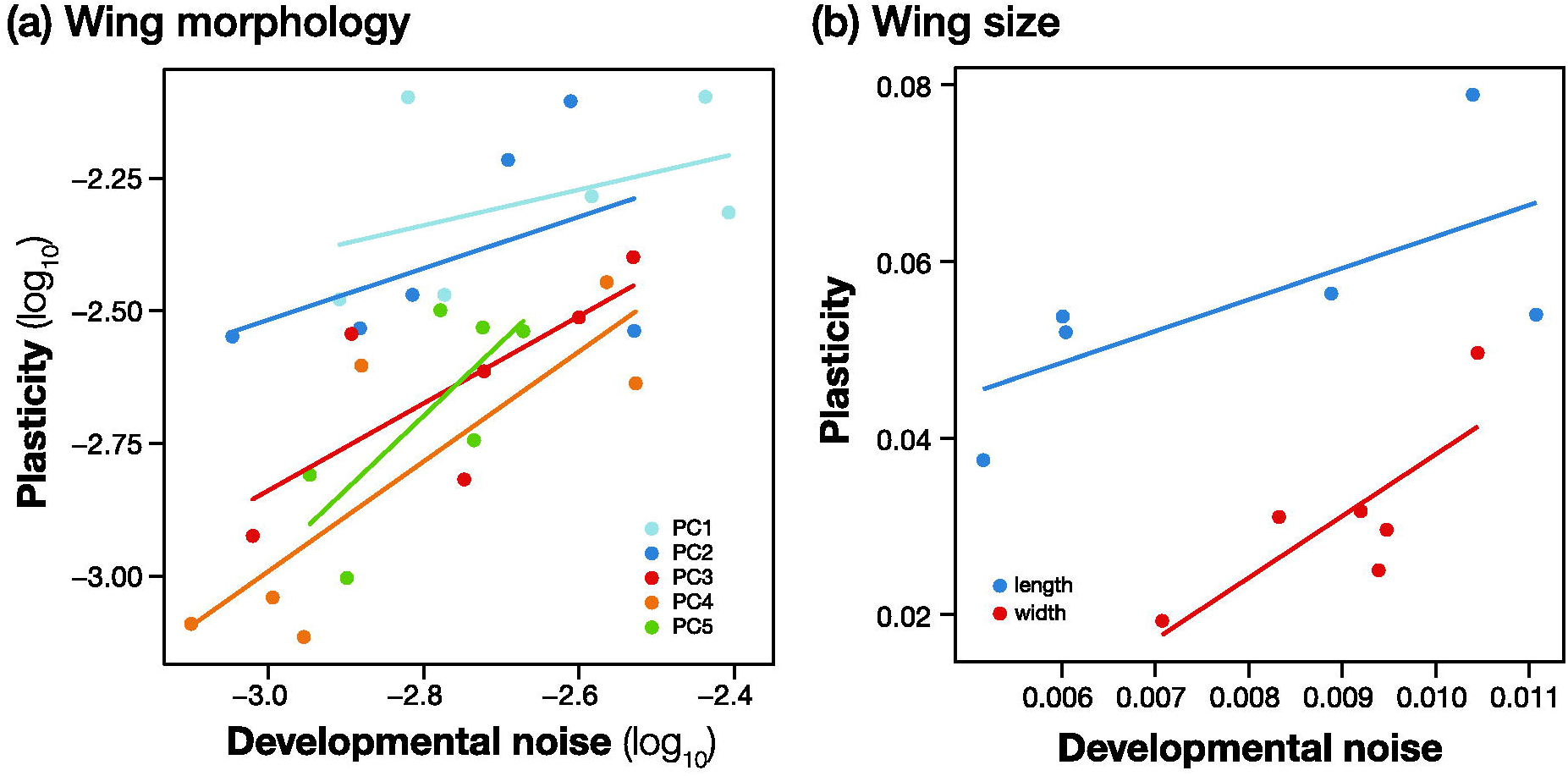
Relationship between the phenotypic plasticity and degree of developmental noise in the wing morphology (a) and wing size (b). Each line represents simple regression lines.

## Discussion

The biological significance of two non-genetic variations, i.e., phenotypic plasticity and developmental noise, in the evolutionary process has become increasingly prominent over the past decades (Furusawa et al., 2005; Price et al., 2003; Rohner et al., 2022; Uller et al., 2018; West-Eberhard, 2003). For example, phenotypic plasticity is suggested to guide adaptive evolution via genetic accommodation (Badyaev, 2011; Levis et al., 2018; Levis & Pfennig, 2016; Pfennig et al., 2010; West-Eberhard, 2003), and the degree of developmental noise among individuals is suggested to be correlated with the rate of evolution (Furusawa et al., 2005; Kaneko & Furusawa, 2006; Rohner et al., 2022; K. Sato et al., 2003). However, these two profoundly different features of non-genetic variation, i.e., phenotypic plasticity and developmental noise, are rarely measured separately, and it remains unclear whether the propensity of a genotype to produce the two types of non-genetic variations are heritable and if they are related. In this study, we quantified the plasticity and developmental noise in the wing morphological traits (size and shape) of *D. simulans* and showed that the wing size and shape responded plastically to different rearing conditions. We also found genetic variation in the propensity for a genotype to produce phenotypic variation through plasticity and developmental noise. Moreover, we demonstrated a positive relationship between phenotypic plasticity and developmental noise. These results add to the increasing body of evidence for the biological significance of non-genetic variation in the evolutionary process. Below, we discuss our three main findings.

Wing morphology, length, and width varied depending on the environment to which the individuals were exposed. In general, the temperate populations of *D. melanogaster* is known to be larger than the tropical populations (David & Capy, 1988; Lack et al., 2016). Accordingly, our results showed that the wings of individuals reared under higher temperature conditions were smaller than those reared under lower temperature conditions. Unlike homeotherms, the growth rate of ectotherms is generally accepted to increase as the temperature increases and drastically decreases after the temperature reaches the maximum growth limiting temperature (Yamahira et al., 2007). Under relatively higher temperature conditions, developmental time becomes shorter due to a high growth rate and shorter developmental times lead to a downsize in wings. This is called the “temperature–size rule” (Atkinson, 1994). Therefore, the downsized wings were thought to be derived from the decreasing developmental time. Interestingly, we found no evidence that the pattern of plastic responses to the three environmental factors (nutrition, light cycle, and temperature) were correlated. We interpret these results as suggesting either that the actual pattern of phenotypic plasticity could not be evaluated by this range of perturbations that we introduced within one environmental factor or that the wing phenotypes can indeed respond differently to different environmental cues.

In wing morphological traits of *D. simulans*, the broad-sense heritability (H^2^) of phenotypic plasticity was 9.4%, and the H^2^ of the developmental noise was 1.4%. These results indicate that there is a propensity for some genotypes to generate more variation in response to environmental cues or local stochastic events during development than others, and these propensities exhibit genetic variance within the fly population. Thus, when selection acts on those propensities either directly or indirectly, they should be able to evolve (Mather, 1953). Our estimates are comparable to the heritability of plasticity reported in morphological traits in *D. melanogaster* (Mackay & Lyman, 2005; Scheiner & Lyman, 1989) but are smaller than those of the behavioral traits, such as chill coma recovery time and startle response (Morgante et al., 2015). Developmental noise showed considerably lower heritability than phenotypic plasticity. Our estimates of H^2^ in the developmental noise were within the range of 0.1%–2%, which are at the lower range of H^2^ or the narrow-sense heritability reported in previous studies (Whitlock & Fowler, 1997). One explanation for these low estimates in wing morphological traits is Fisher’s fundamental theorem of natural selection (Fisher, 1930), which proposes that traits strongly associated with an organism’s fitness will have a lower heritability. Considering the functional significance of wing size and shape for flight performance, it is conceivable that these traits are strongly linked to fitness.

We found two sources of non-genetic variation, i.e., phenotypic plasticity and developmental noise, are correlated. We propose two hypotheses to explain these patterns that are not mutually exclusive. First, a certain gene(s) may govern both the ability to change phenotypic expression in response to external cues (phenotypic plasticity) and through local perturbations during development (developmental noise). Previous studies have suggested that genes representing hubs that stitch together genetic networks are likely to influence the overall phenotypic plasticity or robustness of organisms (Laitinen & Nikoloski, 2019). For example, ELF3 was proposed to be a key hub gene that integrates the developmental and environmental signals in response to temperature (Anwer et al., 2014; Boden et al., 2014; Box et al., 2015). Moreover, HSP-90 is known to contribute to the system robustness or standard genetic variation or developmental noise either directly or indirectly (Mestek Boukhibar & Barkoulas, 2016). Second, both types of non-genetic variation may be produced by the same developmental machinery. Given that the wing morphological traits are likely under strong stabilizing selection, it is conceivable that repeated bouts of selection through the history of fly family *Drosophilidae* have canalized the developmental system to generate adaptive phenotypes more often than the non-adaptive alternatives. This presents the possibility that prior periods of selection have shaped the developmental machinery and determine the pattern of variation in contemporary populations (Jones et al., 2007; Rohner et al., 2022; Uller et al., 2018).

Recently, Rohner and Berger (2023) have shown that developmental noise is correlated with the pattern of genetic and phenotypic variation as well as with the divergence pattern among species (e.g., macroevolution) in wing morphological traits of *Sepsis* flies. Together with previous findings in *Drosophila* flies (Houle et al., 2017) and *Anolis* lizards (McGlothlin et al., 2018), these studies suggest a remarkably consistent and strong correlation between the variational properties across multiple levels of biological organization including evolutionary divergence over tens of millions of years. Our study adds a level to these correlations: the phenotypic plasticity. Interestingly, Rohner and Berger (2023) evaluated the phenotypic plasticity but did not find a correlation between plasticity and developmental noise. We propose two possible explanations for the discrepancy between these results. First, as previously noted, the plastic response in wing morphological traits were small in magnitude in *D. simulans*. The plastic variation in wing shape of *Sepsis* flies may be similarly small, making the relationship challenging to detect. Second, our sample size to estimate the developmental noise is large (n = 87 in Rohner and Berger, 2023, vs. n = 410 in our study), which allowed us to estimate the FA more precisely. The same reasoning could be applied for the other conflicting results regarding the relationship between these two non-genetic variations (Scheiner, 1993). If our arguments are correct, the correlation between plasticity and the other levels of variation should be common, but the detection of this pattern requires precise estimations. There is circumstantial evidence pointing to this possibility (Noble et al., 2019). To fully appreciate the biological significance of these non-genetic variations on the evolutionary process, further studies are required to examine the correlation between them in other organisms and traits.

## Author Contributions

KS and YT conceived and designed the study. KS collected the data. KS analyzed the data, and MT and YT supported for analysis. KS drafted the manuscript. All authors reviewed the manuscript.

## Acknowledgements

The study was funded by a part of KAKENHI (grant no. 20H04857).

## Conflict of Interest

The authors declare no competing interest.

## Data Archiving

All raw data used in this study have been uploaded to the Figshare, 0.6084/m9.figshare.23615622.

## Notes

### Competing Interest Statement

The authors have declared no competing interest.

## References

1. Adams, D. C., & Otárola-Castillo, E. (2013). geomorph: An R package for the collection and analysis of geometric morphometric shape data. Methods in Ecology and Evolution, 4(4), 393–399. https://doi.org/10.1111/2041-210X.12035

2. Anwer, M. U., Boikoglou, E., Herrero, E., Hallstein, M., Davis, A. M., Velikkakam James, G., Nagy, F., & Davis, S. J. (2014). Natural variation reveals that intracellular distribution of ELF3 protein is associated with function in the circadian clock. ELife, 3, e02206. https://doi.org/10.7554/eLife.02206

3. Arnold, S. J. (1992). Constraints on phenotypic evolution. The American Naturalist, 140, 85–107. https://doi.org/10.1086/285398

4. Atkinson, D. (1994). Temperature and organism size-A biological law for ectotherms? Adovances in Ecological Research, 25(1). https://doi.org/10.1016/S0065-2504(08)60212-3

5. Badyaev, A. V. (2011). Origin of the fittest: Link between emergent variation and evolutionary change as a critical question in evolutionary biology. Proceedings of the Royal Society B: Biological Sciences, 278(1714), 1921–1929. https://doi.org/10.1098/rspb.2011.0548

6. Bates, D., Mächler, M., Bolker, B., & Walker, S. (2015). Fitting linear mixed-effects models using lme4. Journal of Statistical Software, 67, 1–48. https://doi.org/10.18637/jss.v067.i01

7. Becker, W. A. (1964). Heritability of a response to an environmental change in chickens. Genetics, 50(5), 783–788. https://doi.org/10.1093%2Fgenetics%2F50.5.783

8. Boden, S. A., Weiss, D., Ross, J. J., Davies, N. W., Trevaskis, B., Chandler, P. M., & Swain, S. M. (2014). EARLY FLOWERING3 regulates flowering in spring barley by mediating gibberellin production and FLOWERING LOCUS T expression. The Plant Cell, 26(4), 1557–1569. https://doi.org/10.1105/tpc.114.123794

9. Box, M. S., Huang, B. E., Domijan, M., Jaeger, K. E., Khattak, A. K., Yoo, S. J., Sedivy, E. L., Jones, D. M., Hearn, T. J., Webb, A. A. R., Grant, A., Locke, J. C. W., & Wigge, P. A. (2015). ELF3 controls thermoresponsive growth in Arabidopsis. Current Biology, 25(2), 194–199. https://doi.org/10.1016/j.cub.2014.10.076

10. Bradshaw, A. D. (1965). Evolutionary significance of phenotypic plasticity in plants. In E. W. Caspari & J. M. Thoday (Eds.), Advances in Genetics (Vol. 13, pp. 115–155). https://doi.org/10.1016/S0065-2660(08)60048-6

11. Burns, I. G. (1992). Influence of plant nutrient concentration on growth rate: Use of a nutrient interruption technique to determine critical concentrations of N, P and K in young plants. Plant and Soil, 142(2), 221–233. https://doi.org/10.1007/BF00010968

12. Careau, V., Wolak, M. E., Carter, P. A., & Garland, T. (2015). Evolution of the additive genetic variance–covariance matrix under continuous directional selection on a complex behavioural phenotype. Proceedings of the Royal Society B: Biological Sciences, 282(1819), 20151119. https://doi.org/10.1098/rspb.2015.1119

13. Danchin, E. (2013). Avatars of information: Towards an inclusive evolutionary synthesis. Trends in Ecology & Evolution, 28(6), 351–358. https://doi.org/10.1016/j.tree.2013.02.010

14. David, J. R., & Capy, P. (1988). Genetic variation of Drosophila melanogaster natural populations. Trends in Genetics, 4(4), 106–111. https://doi.org/10.1016/0168-9525(88)90098-4

15. De Lisle, S. P., Mäenpää, M. I., & Svensson, E. I. (2022). Phenotypic plasticity is aligned with phenological adaptation on both micro-and macroevolutionary timescales. Ecology Letters, 25(4), 790–801. https://doi.org/10.1111/ele.13953

16. Draghi, J. (2020). Developmental noise and ecological opportunity across space can release constraints on the evolution of plasticity. Evolution & Development, 22(1–2), 35–46. https://doi.org/10.1111/ede.12305

17. Fisher, R. A. (1930). The genetical theory of natural selection (pp. 1–302). Clarendon Press. https://doi.org/10.5962/bhl.title.27468

18. Fitzpatrick, M. J., Feder, E., Rowe, L., & Sokolowski, M. B. (2007). Maintaining a behaviour polymorphism by frequency-dependent selection on a single gene. Nature, 447(7141), 210–212. https://doi.org/10.1038/nature05764

19. Furusawa, C., Suzuki, T., Kashiwagi, A., Yomo, T., & Kaneko, K. (2005). Ubiquity of log-normal distributions in intra-cellular reaction dynamics. Biophysics, 1, 25–31. https://doi.org/10.2142/biophysics.1.25

20. Fusco, G., & Minelli, A. (2010). Phenotypic plasticity in development and evolution: Facts and concepts. Philosophical Transactions of the Royal Society B: Biological Sciences, 365(1540), 547–556. https://doi.org/10.1098/rstb.2009.0267

21. Gangestad & Thornhill. (1999). Individual differences in developmental precision and fluctuating asymmetry: A model and its implications. Journal of Evolutionary Biology, 12(2), 402–416. https://doi.org/10.1046/j.1420-9101.1999.00039.x

22. Geiler-Samerotte, K., Bauer, C., Li, S., Ziv, N., Gresham, D., & Siegal, M. (2013). The details in the distributions: Why and how to study phenotypic variability. Current Opinion in Biotechnology, 24(4), 752–759. https://doi.org/10.1016/j.copbio.2013.03.010

23. Ghalambor, C. K., McKAY, J. K., Carroll, S. P., & Reznick, D. N. (2007). Adaptive versus non-adaptive phenotypic plasticity and the potential for contemporary adaptation in new environments. Functional Ecology, 21(3), 394–407. https://doi.org/10.1111/j.1365-2435.2007.01283.x

24. Goswami, A., Binder, W. J., Meachen, J., & O’Keefe, F. R. (2015). The fossil record of phenotypic integration and modularity: A deep-time perspective on developmental and evolutionary dynamics. Proceedings of the National Academy of Sciences, 112(16), 4891–4896. https://doi.org/10.1073/pnas.1403667112

25. Hansen, T. F., & Houle, D. (2008). Measuring and comparing evolvability and constraint in multivariate characters. Journal of Evolutionary Biology, 21(5), 1201–1219. https://doi.org/10.1111/j.1420-9101.2008.01573.x

26. Houle, D. (1997). Comment on “A meta-analysis of the heritability of developmental stability” by Møller and Thornhill. Journal of Evolutionary Biology, 10(1), 17–20. https://doi.org/10.1046/j.1420-9101.1997.10010017.x

27. Houle, D., Bolstad, G. H., van der Linde, K., & Hansen, T. F. (2017). Mutation predicts 40 million years of fly wing evolution. Nature, 548(7668), 447–450. https://doi.org/10.1038/nature23473

28. Hudak, A., & Dybdahl, M. (2023). Phenotypic plasticity under the effects of multiple environmental variables. Evolution, 77(6), 1370–1381. https://doi.org/10.1093/evolut/qpad049

29. Jones, A. G., Arnold, S. J., & Bürger, R. (2007). The mutation matrix and the evolution of evolvability. Evolution, 61(4), 727–745. https://doi.org/10.1111/j.1558-5646.2007.00071.x

30. Kaneko, K., & Furusawa, C. (2006). An evolutionary relationship between genetic variation and phenotypic fluctuation. Journal of Theoretical Biology, 240(1), 78–86. https://doi.org/10.1016/j.jtbi.2005.08.029

31. Kiskowski, M., Glimm, T., Moreno, N., Gamble, T., & Chiari, Y. (2019). Isolating and quantifying the role of developmental noise in generating phenotypic variation. PLOS Computational Biology, 15(4), e1006943. https://doi.org/10.1371/journal.pcbi.1006943

32. Klingenberg, C. P. (2019). Phenotypic plasticity, developmental instability, and robustness: The concepts and how they are connected. Frontiers in Ecology and Evolution, 7, 1–15. https://doi.org/10.3389/fevo.2019.00056

33. Lack, J. B., Yassin, A., Sprengelmeyer, Q. D., Johanning, E. J., David, J. R., & Pool, J. E. (2016). Life history evolution and cellular mechanisms associated with increased size in high altitude Drosophila. Ecology and Evolution, 6(16), 5893–5906. https://doi.org/10.1002/ece3.2327

34. Laitinen, R. A. E., & Nikoloski, Z. (2019). Genetic basis of plasticity in plants. Journal of Experimental Botany, 70(3), 739–745. https://doi.org/10.1093/jxb/ery404

35. Lande, R. (1979). Quantitative genetic analysis of multivariate evolution, applied to brain: Body size allometry. Evolution, 33(1), 402–416. https://doi.org/10.2307/2407630

36. Levis, N. A., Isdaner, A. J., & Pfennig, D. W. (2018). Morphological novelty emerges from pre-existing phenotypic plasticity. Nature Ecology & Evolution, 2(8), 1289–1297. https://doi.org/10.1038/s41559-018-0601-8

37. Levis, N. A., & Pfennig, D. W. (2016). Evaluating ‘plasticity-first’ evolution in nature: Key criteria and empirical approaches. Trends in Ecology & Evolution, 31(7), 563–574. https://doi.org/10.1016/j.tree.2016.03.012

38. Lindestad, O., Wheat, C. W., Nylin, S., & Gotthard, K. (2019). Local adaptation of photoperiodic plasticity maintains life cycle variation within latitudes in a butterfly. Ecology, 100(1), e02550. https://doi.org/10.1002/ecy.2550

39. Mackay, T. F. C., & Lyman, R. F. (2005). Drosophila bristles and the nature of quantitative genetic variation. Philosophical Transactions of the Royal Society B: Biological Sciences, 360(1459), 1513–1527. https://doi.org/10.1098/rstb.2005.1672

40. Matesanz, S., Horgan-Kobelski, T., & Sultan, S. E. (2012). Phenotypic plasticity and population differentiation in an ongoing species invasion. PloS One, 7(9), e44955. https://doi.org/10.1371/journal.pone.0044955

41. Mather, K. (1953). Genetical control of stability in development. Heredity, 7(3), 297–336. https://doi.org/10.1038/hdy.1953.41

42. McGlothlin, J. W., Kobiela, M. E., Wright, H. V., Mahler, D. L., Kolbe, J. J., Losos, J. B., & Brodie, E. D. (2018). Adaptive radiation along a deeply conserved genetic line of least resistance in Anolis lizards. Evolution Letters, 2(4), 310–322. https://doi.org/10.1002/evl3.72

43. Mestek Boukhibar, L., & Barkoulas, M. (2016). The developmental genetics of biological robustness. Annals of Botany, 117(5), 699–707. https://doi.org/10.1093/aob/mcv128

44. Morgante, F., Sørensen, P., Sorensen, D. A., Maltecca, C., & Mackay, T. F. C. (2015). Genetic architecture of micro-environmental plasticity in Drosophila melanogaster. Scientific Reports, 5(1), 9785. https://doi.org/10.1038/srep09785

45. Noble, D. W. A., Radersma, R., & Uller, T. (2019). Plastic responses to novel environments are biased towards phenotype dimensions with high additive genetic variation. Proceedings of the National Academy of Sciences, 116(27), 13452–13461. https://doi.org/10.1073/pnas.1821066116

46. Nyamukondiwa, C., Terblanche, J. S., Marshall, K. E., & Sinclair, B. J. (2011). Basal cold but not heat tolerance constrains plasticity among Drosophila species (Diptera: Drosophilidae). Journal of Evolutionary Biology, 24(9), 1927–1938. https://doi.org/10.1111/j.1420-9101.2011.02324.x

47. Palmer, A. R., & Strobeck, C. (1986). Fluctuating asymmetry: Measurement, analysis, patterns. Annual Review of Ecology and Systematics, 17, 391–421. https://doi.org/10.1146/ANNUREV.ES.17.110186.002135

48. Pfennig, D. W., Wund, M. A., Snell-Rood, E. C., Cruickshank, T., Schlichting, C. D., & Moczek, A. P. (2010). Phenotypic plasticity’s impacts on diversification and speciation. Trends in Ecology & Evolution, 25(8), 459–467. https://doi.org/10.1016/j.tree.2010.05.006

49. Pigliucci, M., Murren, C. J., & Schlichting, C. D. (2006). Phenotypic plasticity and evolution by genetic assimilation. Journal of Experimental Biology, 209(12), 2362–2367. https://doi.org/10.1242/jeb.02070

50. Porto, A., & Voje, K. L. (2020). ML-morph: A fast, accurate and general approach for automated detection and landmarking of biological structures in images. Methods in Ecology and Evolution, 11(4), 500–512. https://doi.org/10.1111/2041-210X.13373

51. Price, T. D., Qvarnström, A., & Irwin, D. E. (2003). The role of phenotypic plasticity in driving genetic evolution. Proceedings of the Royal Society B: Biological Sciences, 270(1523), 1433–1440. https://doi.org/10.1098/rspb.2003.2372

52. Rohner, P. T., & Berger, D. (2023). Developmental bias predicts 60 million years of wing shape evolution. Proceedings of the National Academy of Sciences, 120(19), e2211210120. https://doi.org/10.1073/pnas.2211210120

53. Rohner, P. T., Hu, Y., & Moczek, A. P. (2022). Developmental bias in the evolution and plasticity of beetle horn shape. Proceedings of the Royal Society B: Biological Sciences, 289(1983), 20221441. https://doi.org/10.1098/rspb.2022.1441

54. Sato, A., & Takahashi, Y. (2022). Responses in thermal tolerance and daily activity rhythm to urban stress in Drosophila suzukii. Ecology and Evolution, 12(12), e9616. https://doi.org/10.1002/ece3.9616

55. Sato, K., Ito, Y., Yomo, T., & Kaneko, K. (2003). On the relation between fluctuation and response in biological systems. Proceedings of the National Academy of Sciences, 100(24), 14086–14090. https://doi.org/10.1073/pnas.2334996100

56. Scheiner, S. M. (1993). Genetics and evolution of phenotypic plasticity. Annual Review of Ecology and Systematics, 24(1), 35–68. https://doi.org/10.1146/annurev.es.24.110193.000343

57. Scheiner, S. M., Caplan, R. L., & Lyman, R. F. (1991). The genetics of phenotypic plasticity. III. Genetic correlations and fluctuating asymmetries. Journal of Evolutionary Biology, 4(1), 51–68. https://doi.org/10.1046/j.1420-9101.1991.4010051.x

58. Scheiner, S. M., & Lyman, R. F. (1989). The genetics of phenotypic plasticity I. Heritability. Journal of Evolutionary Biology, 2(2), 95–107. https://doi.org/10.1046/j.1420-9101.1989.2020095.x

59. Schluter, D. (1996). Adaptive radiation along genetic lines of least resistance. Evolution, 50(5), 1766–1774. https://doi.org/10.2307/2410734

60. Shehata, T. E., & Marr, A. G. (1971). Effect of nutrient concentration on the growth of Escherichia coli. Journal of Bacteriology, 107(1), 210–216. https://doi.org/10.1128%2Fjb.107.1.210-216.1971

61. Uller, T., Moczek, A. P., Watson, R. A., Brakefield, P. M., & Laland, K. N. (2018). Developmental bias and evolution: A regulatory network perspective. Genetics, 209(4), 949–966. https://doi.org/10.1534/genetics.118.300995

62. Van Valen, L. (1962). A study of fluctuating asymmetry. Evolution, 16(2), 125–142. https://doi.org/10.2307/2406192

63. West-Eberhard, M. J. (2003). Developmental Plasticity and Evolution. Oxford University Press.

64. Westneat, David. F., Potts, L. J., Sasser, K. L., & Shaffer, J. D. (2019). Causes and consequences of phenotypic plasticity in complex environments. Trends in Ecology & Evolution, 34(6), 555–568. https://doi.org/10.1016/j.tree.2019.02.010

65. Whitlock, M. C., & Fowler, K. (1997). The instability of studies of instability. Journal of Evolutionary Biology, 10(1), 63–67. https://doi.org/10.1046/j.1420-9101.1997.10010063.x

66. Wilson, A. J., Réale, D., Clements, M. N., Morrissey, M. M., Postma, E., Walling, C. A., Kruuk, L. E. B., & Nussey, D. H. (2010). An ecologist’s guide to the animal model. Journal of Animal Ecology, 79(1), 13–26. https://doi.org/10.1111/j.1365-2656.2009.01639.x

67. Yamahira, K., Kawajiri, M., Takeshi, K., & Irie, T. (2007). Inter-and intrapopulation variation in thermal reaction norms for growth rate: Evolution of latitudinal compensation in ectotherms with a genetic constraint. Evolution, 61(7), 1577–1589. https://doi.org/10.1111/j.1558-5646.2007.00130.x

