## Supplementary material for "Developmental noise and phenotypic plasticity are correlated in *Drosophila simulans*": SI

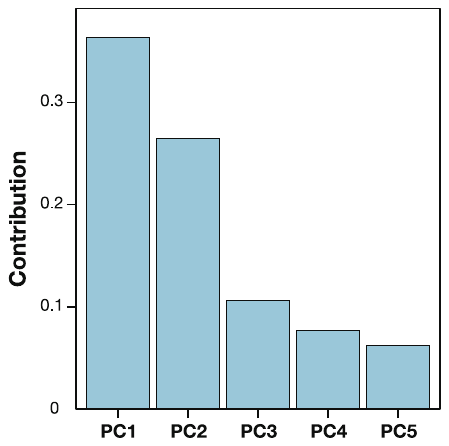

Figure S1. Top five principal component axes whose contribution was more than 5%.
